# Evolutionary history of the global emergence of the *Escherichia coli* epidemic clone ST131

**DOI:** 10.1101/030668

**Authors:** Nicole Stoesser, Anna E. Sheppard, Louise Pankhurst, Nicola de Maio, Catrin E. Moore, Robert Sebra, Paul Turner, Luke W. Anson, Andrew Kasarskis, Elizabeth M. Batty, Veronica Kos, Daniel J. Wilson, Rattanaphone Phetsouvanh, David Wyllie, Evgeni Sokurenko, Amee R. Manges, Timothy J. Johnson, Lance B. Price, Timothy E. A. Peto, James R. Johnson, Xavier Didelot, A. Sarah Walker, Derrick W. Crook, Modernising Medical Microbiology Informatics Group (MMMIG)

## Abstract

**Background:** *Escherichia coli* sequence type 131 (ST131) has emerged globally as the most predominant lineage within this clinically important species, and its association with fluoroquinolone and extended-spectrum cephalosporin resistance impacts significantly on treatment. The evolutionary histories of this lineage, and of important antimicrobial resistance elements within it, remain unclearly defined.

**Results:** This study of the largest worldwide collection (n = 215) of sequenced ST131 *E. coli* isolates to date demonstrates that clonal expansion of two previously recognized antimicrobial-resistant clades, C1/*H*30R and C2/*H*30Rx, started around 25 years ago, consistent with the widespread introduction of fluoroquinolones and extended-spectrum cephalosporins in clinical medicine. These two clades appear to have emerged in the United States, with the expansion of the C2/*H*30Rx clade driven by the acquisition of a *bla*_CTX-M-15_-containing IncFII-like plasmid that has subsequently undergone extensive rearrangement. Several other evolutionary processes influencing the trajectory of this drug-resistant lineage are described, including sporadic acquisitions of CTX-M resistance plasmids, and chromosomal integration of *bla*_CTX-M_ within sub-clusters followed by vertical evolution. These processes are also occurring for another family of CTX-M gene variants more recently observed amongst ST131, the *bla***_C_**_TX-M-14/14-like_ group.

**Conclusions:** The complexity of the evolutionary history of ST131 has important implications for antimicrobial resistance surveillance, epidemiological analysis, and control of emerging clinical lineages of *E. coli.* These data also highlight the global imperative to reduce specific antibiotic selection pressures, and demonstrate the important and varied roles played by plasmids and other mobile genetic elements in the perpetuation of antimicrobial resistance within lineages.

## IMPORTANCE

*Escherichia coli,* perennially a major bacterial pathogen, is becoming increasingly difficult to manage due to emerging resistance to all preferred antimicrobials. Resistance is concentrated within specific *E. coli* lineages, such as sequence type (ST) 131. Clarification of the genetic basis for clonally-associated resistance is key to devising intervention strategies.

We used high-resolution genomic analysis of a large global collection of ST131 isolates to define the evolutionary history of extended-spectrum beta-lactamase production in ST131. We documented diverse contributory genetic processes, including stable chromosomal integrations of resistance genes, persistence and evolution of mobile resistance elements within sub-lineages, and sporadic acquisition of different resistance elements. Both global distribution and regional segregation were evident. The diversity of resistance element acquisition and propagation within ST131 indicates aneed for flexible approaches to control and for ongoing surveillance.

## BACKGROUND

Resistance to extended-spectrum cephalosporins in extra-intestinal pathogenic *Escherichia coli* (ExPEC) represents a major clinical challenge and is commonly caused by the presence of extended-spectrum beta-lactamases (ESBLs). Most ESBL-associated *E. coli* infections are due to a recently emerged, globally distributed ExPEC clone, sequence type (ST) 131. ST131 corresponds to serogroup 025b [1, 2] and belongs to phylogenetic group B2 [3, 4], It remains unclear which features of this clone have resulted in its recent widespread clinical dominance, although antimicrobial resistance and virulence factors are suspected contributors [5].

The *bla*_CTX-M-15_ beta-lactamasegene is the dominant ESBL gene in ST131, but other genetically divergent CTX-M genes also occur in this ST, particularly *bla***_C_**_TX-M-14/14-like_ variants, e.g. in Canada, China, and Spain [6, 7], The almost contemporaneous identification of *bla*_CTX-M_ in ST131 strains from multiple geographic locations suggests repeated acquisition via multiple horizontal gene transfer events [8], Consistently, both *bla*_CTX-M-15_ and *bla*_CTX-M-14/14-like_ variants occur on conjugative plasmids, especially multi-replicon IncFII plasmids additionally harboring FIA/FIB replicons [9].

Other data, however, suggest that the widespread distribution of these genes is mediated by clonal expansion of CTX-M-containing strains and global dissemination [10]. This is also a plausible hypothesis, since CTX-M plasmids can be inherited stably and *bla*_CTX-M-15_ and *bla*_CTX-M-14_ variants can also integrate into the chromosome [11-13], Nevertheless, clonal expansion of *E. coli* strains with chromosomally integrated *bla*_CTX-M_ has not yet been demonstrated.

Two recent studies used whole genome sequence (WGS) data to investigate the population structure of ST131. The first found that ST131 expansion in the US has been driven by a single sub-lineage, *H*30, defined by the presence of a specific fimbrial adhesin allele, *fimH*30. Within *H*30, nested clades have emerged: *H*30R, containing mutations in the chromosomal genes *gyrA* and *parC* that confer fluoroquinolone resistance, and *H*30Rx, containing the same *gyrA* and *parC* mutations but additionally associated with *bla*_CTX-M-15_ [11]. The second study [14], which included samples from six locations around the world, resolved the ST131 population structure into three clades, A, B, and C, with clade C comprising two sub-groups, C1 and C2, corresponding to the *H*30R and *H*30Rx clades. However, this study included only four isolates from Asia, where ESBL ExPEC prevalence may be highest [15], Furthermore, neither study directly tested the competing hypotheses that ESBL dissemination in ST131 has occurred through multiple horizontal gene transfer events versus clonal expansion.

Here we used a broader set of ST131 WGS data, including many more isolates from Asia [16] and CTX-M-14/14-like-containing strains, alongside a subset of CTX-M plasmid sequences, to estimate the contribution of each potential route of dissemination to the worldwide prevalence ofST131.

## RESULTS

The 215 ST131 genome sequences analyzed included 67 strains from various locations in Southeast Asia, 33 from Oxford in the United Kingdom, 11 from a global resistance surveillance program at AstraZeneca, 8 from Canada and 96 predominantly North American isolates previously reported by Price et al [11] (details on new isolates in Supplementary Table S1; these strains included both human and animal, and clinical and carriage isolates.)

### Asian ST131 strains are consistent with the previously described core phylogeny, and the C1/*H*30 and C2/*H*30Rx clades emerged from a North American ancestor

For the 4,717,338 sites in the SE15 ST131 reference genome [17], the mean mapping call rate across the dataset was 93.3%. In total, 40,057 (0.85%) sites were variable, with 6,879 (0.15%) representing core, single nucleotide variants (SNVs) called in all 215 isolates. Overall, 611,770 (13%) sites were in recombinant regions, including 4,120 core SNVs, leaving 2,759 core, non-recombinant SNVs for phylogenetic analysis.

Consistent with the two previous WGS-based ST131 phylogenies [11, 14], the time-scaled phylogeny inferred from this ST131 dataset (which included >10 times more Asian isolates than considered previously), comprised three clades (Fig. 1),A(n = 25), B (n = 51), and C(n = 139), with C containing two sub-clades, C1 (n = 57) and C2 (n = 82), characterized by the presence vs. absence respectively of *bla*_CTX-M-15_ [14]. B and C1 were in fact paraphyletic groups rather than monophyletic clades, but we followed previous notation by calling them clades nonetheless. Isolates from all geographic regions were identified within each clade, although there were smaller, geographically restricted clusters within these (Fig. 1, tip color). This supports both global transmission and localized clonal expansion following specific introductions into a geographic locality.

**Figure 1.**
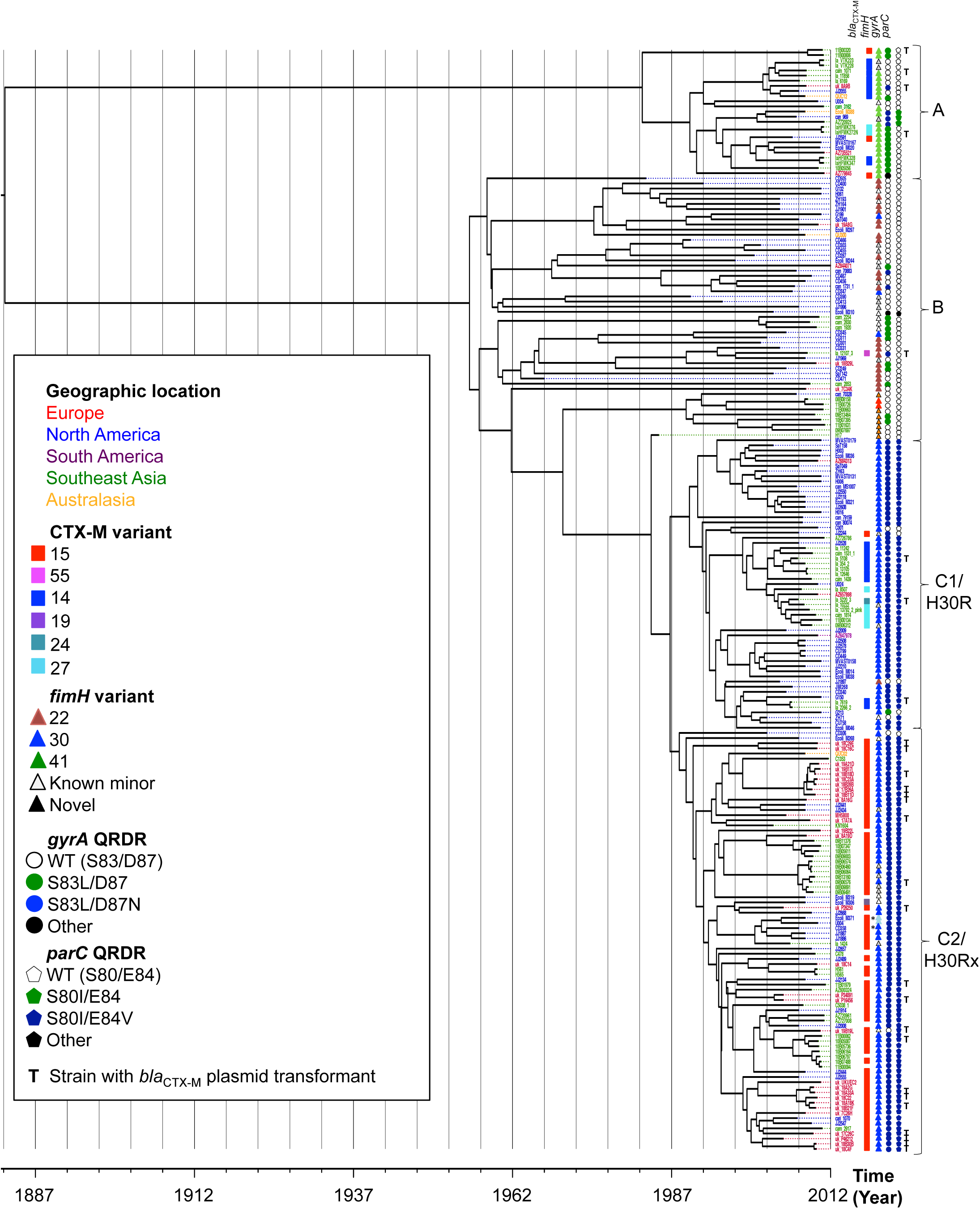
Time-scaled phylogeny of ST131 *E. coli* (n = 215), with associated *bla*_CTX-M_ */fimH* variants, and quinolone-resistance determining region (QRDR) mutations in *gyrA* (WT = wild type QRDR). Curly brackets represent ST131 clades as described in the text. Tips are colored by geographic region, as per key. **T** denotes *bla*_CTX-M_ plasmid transformant generated for strain; * denotes cases with putative deletions in the assembled *bla*_CTX-M-15_ gene.

The estimated time to most recent common ancestor (TMRCA) for the whole genomic dataset was ~130 years ago, when clade A diverged from clades B and C. Twenty-five years ago, clade C emerged out of the paraphyletic clade B, which was quickly followed by the split between sub-clades C1 and C2. The number of core SNVs separating the clades was approximately 250 for clades A vs. B/C, 50-60 for clades B vs. C1/C2, and 10-30 for clades C1 vs. C2. The evolutionary rate of ST131 was estimated in BEAST (see methods) at 2.46×10^−7^ mutations per site per year (95% CI: 2.18-2.75×10^−7^), equating to 1.00 (95% CI: 0.89-1.12) mutation per genome per year.

All possible geographic origins of the root of the ST131 lineage were inferred to be equally likely since the root is far back in time relative to the estimated migration rates. Clade A was inferred to originate in Southeast Asia with ~70% confidence (78% when the unsampled deme was included in the model – see methods), and the B/C clades from North America with ~88% confidence. The ancestral origin of C1/*H*30 and C2/*H*30Rx was strongly inferred as being in North America (98% confidence; 85% confidence when the unsampled deme was included in the model) with subsequent dissemination to Europe and Asia. Locations of more recent nodes are inferred with high confidence, as expected [18].

### *bla*_CTX-M_, *fimH* and *gyrA* variants are strongly associated with specific ST131 clades

Overall, 105 (49%) ST131 isolates harbored *bla*_CTX_*_-_*_M_: *bla*_CTX-M-15_ was found in 74 isolates (34%), *bla*_CTX-M-14_ in 20 (9%), *bla*_CTX-M-27_ in 8 (4%), with one isolate with each of *bla*_CTX-M-19_, *bla*_CTX-M-24_, and *bla*_CTX-M-55_ (Fig. 1). *bla*_CTX-M-15_ was almost completely restricted to the C2 clade, as described previously[14], occurring in 69/82 (84%) C2 isolates but only sporadically in other clades (4/133 ;p < 0.001, Fisher’s exact test). *bla*_CTX-M-14_ and *bla*_CTX-M-27_ were also clustered within the two different clades A and C1, and completely absent from B and C2 (Fig. 1). Overall, the presence of shared *bla*_CTX-M_ variants within clusters was constrained to those with a TMRCA of less than 25 years, suggestive of the emergence of *bla*_CTX-M_ within ST131 after the widespread introduction of third generation cephalosporins in clinical practice.

The most common *fimH* variant was *fimH*30 (n = 123; 57%), followed by *fimH*22 (n = 24; 11%) and *fimH*41 (n = 21; 10%), whereas 23 isolates had novel *fimH* variants, and one was *fimH*-null. As observed for bla_CTX-M_*, fimH* alleles were strongly associated with clade, with 21/25 (84%) isolates in clade A having *fimH*41, 23/51 (45%) in clade B having *fimH*22, and 122/139 (88%) in clade C having *fimH*30 (p < 0.001; Fisher’s exact test).

Fluoroquinolone resistance mutations in *gyrA* and *parC* were also clade-associated, with isolates in clades A and B typically having no or only single mutations in these genes’ quinolone-resistance determining regions (QRDR) (Fig. 1). In contrast, most clade C isolates had double mutations in both *gyrA* and *parC*, shown to confer high-level fluoroquinolone resistance [19] (132/139 [95%]). The seven clade C isolates without these mutations (5 in C1 and 2 in C2) were sporadic, with two having non-*fimH*30 variants, suggesting intermittent recombination events affecting *gyrA, parC,* and *fimH.* The emergence of this double mutation, high-level fluoroquinolone-resistant genotype dated to 25-40 years ago, consistent with the introduction of fluoroquinolones in clinical practice.

### *bla*_CTX-M_**_-_**_15_ in clade C2 is present in a consistent but short flanking structure, frequently truncated by IS*26* elements and within different genetic backgrounds

In four of the 74 *bla*_CTX-M-15_-containing isolates, *bla*_CTX-M-15_ was present on two different contigs (C1353, JJ2643, CD358, JJ2434). In another isolate (JJ2547), the assembled contig with *bla*_CTX-M-15_ contained a series of “N”s, suggesting possible uncertainty around the contig assembly or multiple locations of the gene. These five isolates were excluded from further analysis of flanking regions. In the 69 remaining isolates (3 in A, 1 in C1, 65 in C2), *bla*_CTX-M-15_ was found downstream of a homologous tract of 48bp preceded by an IS*Ecp1* right-end inverted repeat region (IRR-R), and upstream of a homologous tract of 46bp followed by ORF477. This is consistent with the introduction of an IS*Ecp1-bla*_CTX_*_-_*_M-15_-ORF477 unit within ST131, and subsequent rearrangement events affecting this structure.

In clade C2, *bla*_CTX-M-15_ was integrated into the chromosome of 8/65 (12%) isolates, with four unique integration events, one of which was stably present in a sub-cluster of five isolates with TMRCA in 2002 and spread across two geographical regions (Fig. 2). All chromosomal integration events were associated with an intact IS*Ecp1* upstream of *bla*_CTX-M-15_. In three isolates the IS*Ecp1-bla*_CTX_**_-_**_M-15_**-**ORF477 unit was flanked by 5bp target site duplications consistent with transposition, and in one isolate the ORF477 was truncated, suggestive of either one-ended transposition [20], or standard transposition followed by a deletion event (Fig. 2).

**Figure 2.**
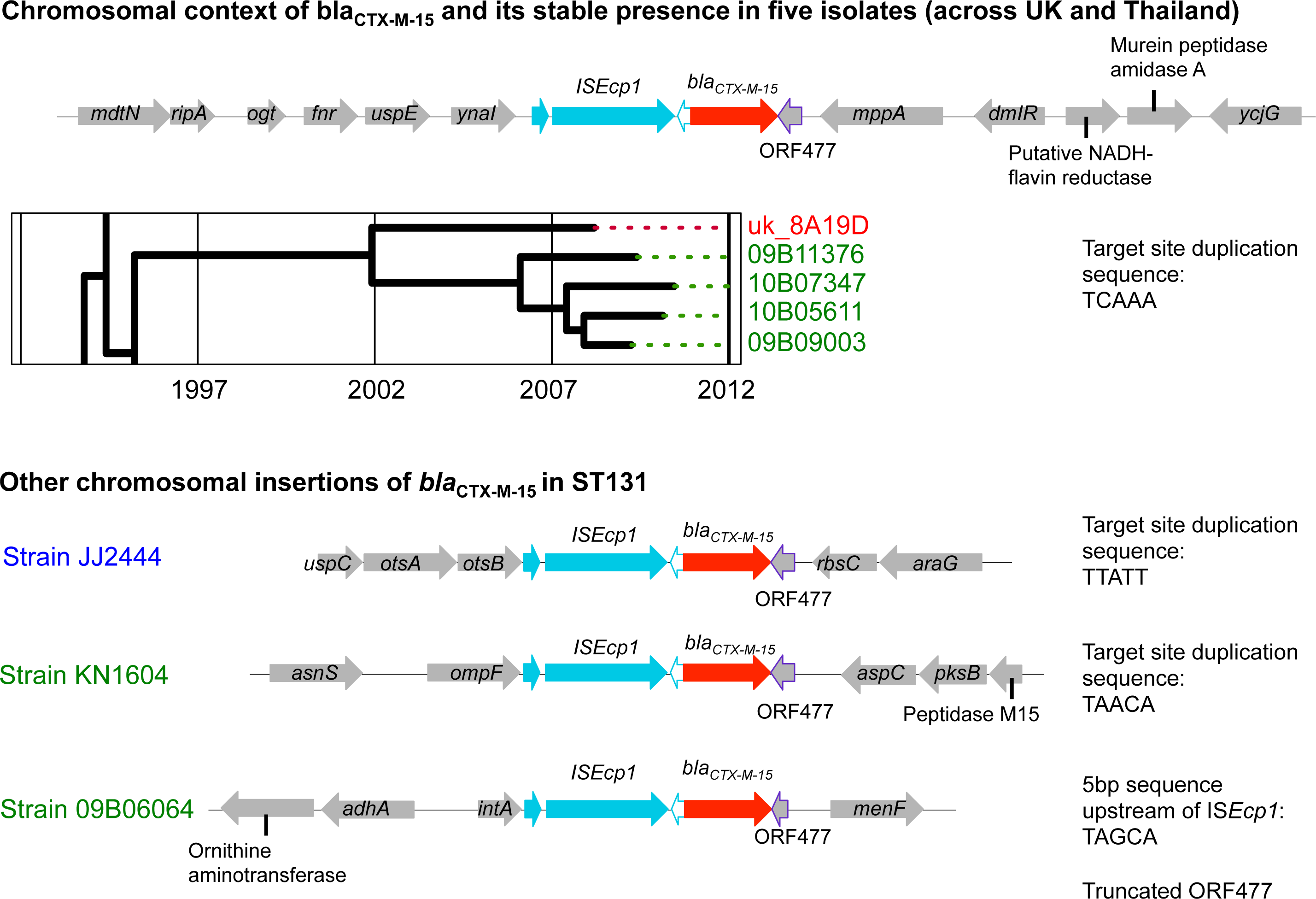
Chromosomal location of IS*Ecp1*-mediated *bla*_CTX-M-15_ insertion events, and evidence of acquisition and evolution by descent (inset phylogeny) over approximately 8 years across two geographic regions (Oxford, UK; Mae Sot, Thai-Myanmar border). Coloring of isolate names represents geographic location (red = Europe, green = South-East Asia, blue = North America).

In 27 of the 57 remaining C2 isolates, *bla*_CTX-M-15_ appeared plasmid-associated, either present in plasmid transformants (n = 20) or flanked by likely plasmid-associated sequences in the contig assemblies (n = 7). In the remaining 30 isolates the location of *bla*_CTX-M-15_ could not be defined due to limitations of the short-read assemblies. In all 57 isolates the upstream sequence was either an intact or truncated IS*Ecp1* sequence, and in 51/57 (89%) isolates the sequence downstream of ORF477 was either an intact or truncated IS*SWi1*-like (Tn*2*-like) structure (Fig. 3). In 12 isolates distributed throughout clade C2, a continuation of the IS*SWil*-like sequence was also observed upstream of the IS*Ecp1* sequence, consistent with the IS*Ecp1* element (flanked by a pair of 5bp repeats, all TCATA) being nested within a complete or partial IS*SWil*-like transposon. In 40/57 (65%) isolates IS*26* repeat regions truncated either or both of these upstream and downstream contexts (Fig. 3).

**Figure 3.**
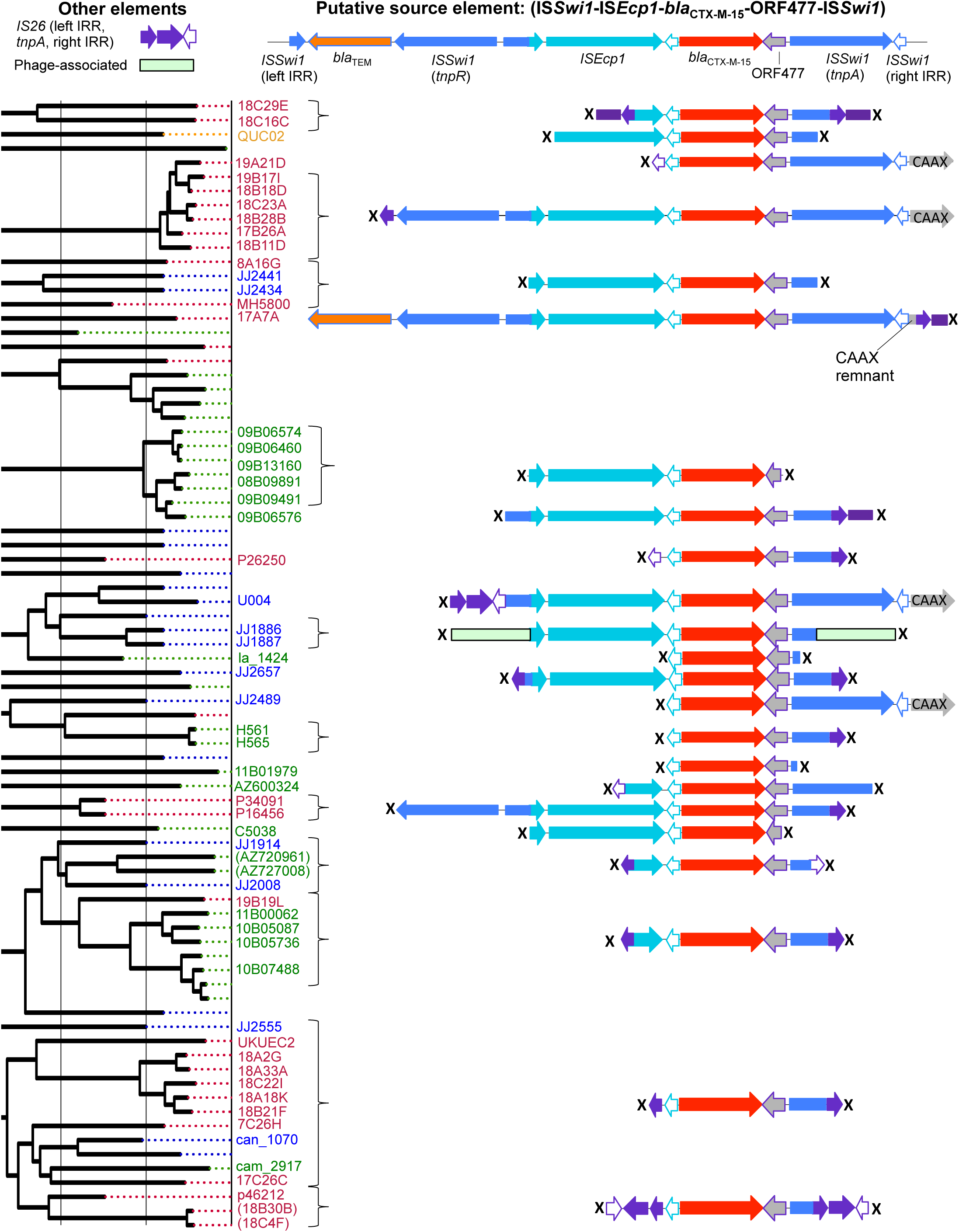
Genetic flanking context of *bla*_CTX-M-15_ region for clade C2/H30Rx isolates. Top: putative source element. Left: C2/H30Rx phylogeny. Many contexts are limited by the extent of the assembled region around the *bla*_CTX-M-15_ gene (marked with “X”). For all similarly colored, vertically aligned regions below the “Putative source element”, sequence identity is 100%. Curly brackets cluster those isolates with identical flanking sequences. Flanking contexts are not shown and tip labels are omitted for isolates with known chromosomal integration of *bla*_CTX-M_, for *bla*_CTX-M_ negative isolates in the clade, or for isolates where the flanking sequence was not evaluable (see “Results”).

### *bla*_CTX-M-14_ and *bla*_CTX-M-27_ are present in diverse genetic backgrounds, and within a common IS*Ecp1-*IS*903B* transposition unit

For *bla*_CTX-M-14_, evidence of chromosomal integration and propagation by descent was also found: two related *bla*_CTX-M-14_ isolates had IS*Ecp1*-mediated chromosomal integration of *bla*_CTX-M-14_ downstream of the *gatY* gene (clade A, isolates HFMK328 and HFMK347). Six isolates had plasmid-associated *bla*_CTX-M-14_ based on annotated flanking sequences/transformants, whereas for the rest (n = 14) the location of *bla*_CTX-M-14_ was uncertain due to limitations of the *de novo* assemblies.

In all these isolates IS*Ecp1* was consistently located upstream of *bla*_CTX-M-14_, as with *bla*_CTX-M-15_, but at only 43bp distance, and the downstream flanking sequences were composed of either intact or truncated IS*903B* elements. In clade A, the genetic flanking sequences surrounding *bla*_CTX-M-14_ were consistent with the host strain sub-cluster, and homologous over the observed contig length within this sub-cluster (Supplementary Figure S1, Clade A CTX-M-14 sub-cluster [i]), supporting a single *bla*_CTX-M-14_ plasmid acquisition event followed by either evolution with plasmid inheritance or subsequent transfer of a *bla*_CTX-M-14_-containing genetic unit within the sub-cluster. The flanking sequence for isolate la_5108_T in clade Cl also incorporated an IS*Ec23* element downstream of IS*903B*, and was homologous to that in clade A CTX-M-14 sub-cluster [i]), (Supplementary Figure S2), suggesting horizontal transfer of this genetic unit between clades.

Six of eight isolates with *bla*_CTX-M-27_ were closely related in clade C1, again supporting a single plasmid acquisition event. However, they also all contained bilateral truncation of the IS*Ecp1*-*bla*_CTX-M_-IS*903B* structure by IS*26* elements, which occurred in four different contexts, suggesting frequent IS*26*-mediated *bla*_CTX-M-27_ transposition events within this sub-cluster (Supplementary Figure S2).

### Plasmid replicon analysis demonstrates a degree of clade-associated plasmid segregation suggestive of ancient IncF plasmid acquisition events

The predominant replicon family was IncF, identified in 206/215 ST131 isolates (96%). Specific IncF variants differed in frequency, with FII found in 199/215 isolates (93%), FIB in 155/215 (72%), FIA in 145/215 (67%), and FIC in 17/215 (8%). Specific IncF replicons and combinations thereof were clade-associated (Table 1). A number of non-F Inc types were also identified; of these, IncH was associated with clade B and Incl with clade C1 (Table 1). Col-like plasmids were also common (189/215 isolates [88%]); however, there was no clear association of any Col-type with clade (Supplementary Figure S3).

**Table 1.**
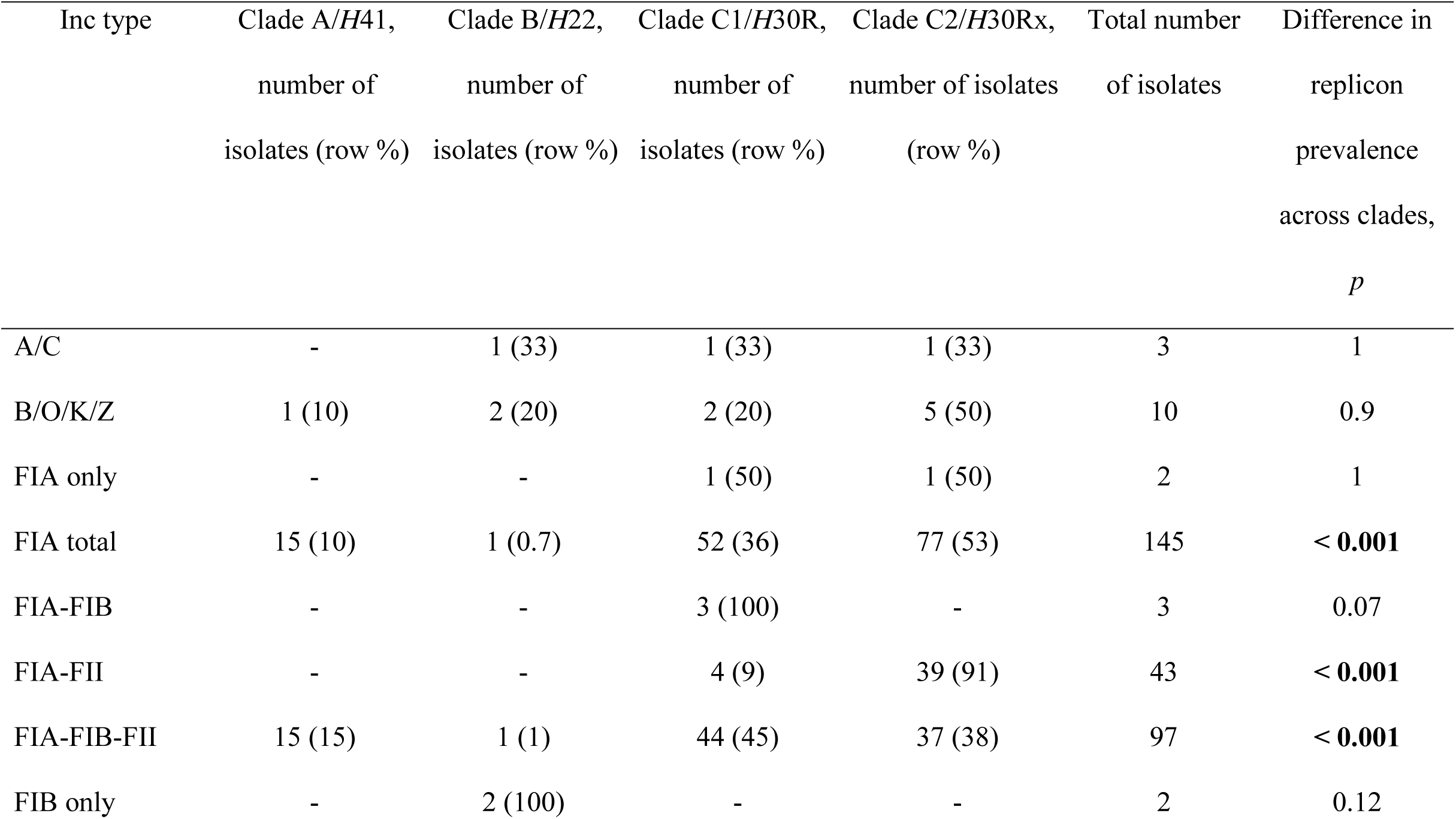
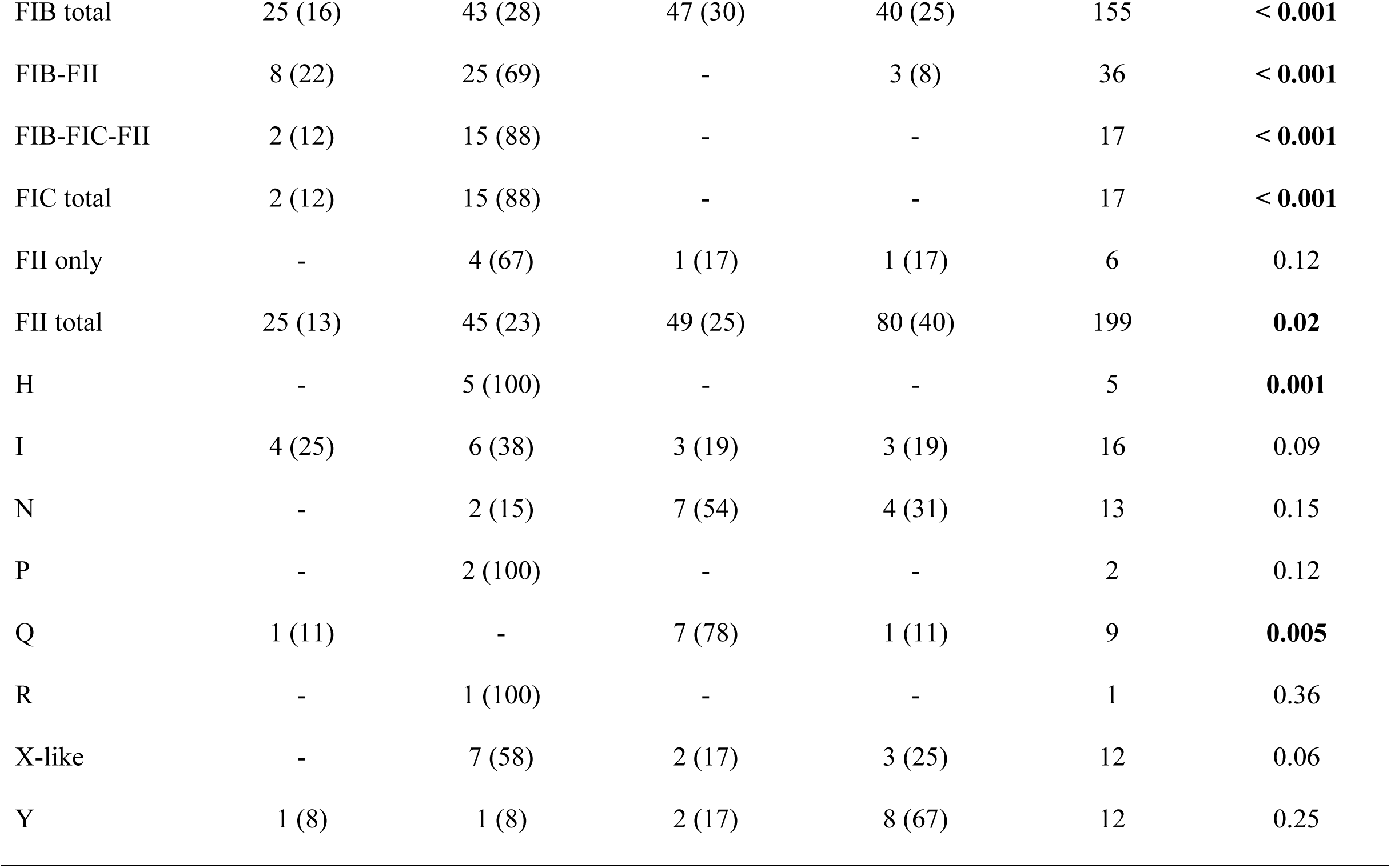
Plasmid replicon families/types by clade

A specific FII variant (GenBank: AY458016; pC15-1a; consistent with pMLST allele 2) was significantly associated with clade C2 (48/82 C2 isolates versus 18/153 non-C2 isolates, p < 0.001, Fisher’s exact test). Within clade C2 a further 23 isolates had eight different FII_AY458016-like variants containing up to 12 SNVs between them; almost all of these variants were in isolates with FIA-FIB-FII replicon combinations (Supplementary Figure S3). Of the 11 clade C2 isolates without FII_AY458016-like replicon variants, four contained a plasmid with a different FII replicon (GenBank: AJ851089; pRSB107, 35 SNVs different from FII_AY458016; consistent with pMLST allele 1), five had chromosomally integrated *bla*_CTX-M-15_ (of which four also contained an FII_AJ851089-like plasmid), one was *bla*_CTX-M_ negative, and one contained deletions in *bla*_CTX-M-15_. There were only nine clade C2 isolates with FII_AY458016-like replicons but no *bla*_CTX-M-15_. The different FII replicon containing *bla*_CTX-M-15_ in clade C2, FII_AJ851089, was also clade-associated, being found predominantly in clades A (13/25 isolates, 52%) and C1 (41/57, 72%) rather than B (12/51, 24%) and C2 (8/82, 10%) (p < 0.0001, Fisher’s exacttest) (Supplementary Figure S3). Overall, this strongly suggests the ancestral acquisition of the FII_AY458016 replicon within clade C2, its association with *bla*_CTX-M-15_ and the expansion of the clade, its evolution in the presence ofFIA-FIB, and its sporadic loss.

### Plasmid transformants demonstrate similarities and differences in *bla*_CTX-M-15_ plasmids from ST131 clades and other sequence types

Sequence data were generated for 30 transformed *bla*_CTX-M_ plasmids (relevant source strains labeled “T” on Fig. 1): four from clade A, containing *bla*_CTX-M-15_ (n = 1), *bla*_CTX-M-14_ (n = 2), and *bla*_CTX-M-27_ (n = 1); one from clade B, containing *bla*_CTX-M-55_; three from clade C1, containing *bla*_CTX-M-14_ (n = 2) and *bla*_CTX-M-24_ (n = 1); 20 from clade C2, containing *bla*_CTX-M-15_; and two *bla*_CTX-M-15_ plasmids from non-ST131 isolates (Supplementary Table S2). A comparison of the mean percentage pairwise differences between all transformant pairs versus the divergence time of the two host strains demonstrated that all *bla*_CTX-M_ transformant plasmids shared at least 10% homology but could be genetically divergent (Fig. 4); plasmids found in different STs could be very similar (up to ~90% sequence homology); and plasmid genetic similarity correlated with host strain divergence time for recently diverged host strains (up to ~30 years) but was much more variable for more remotely diverged host strains.

**Figure 4.**
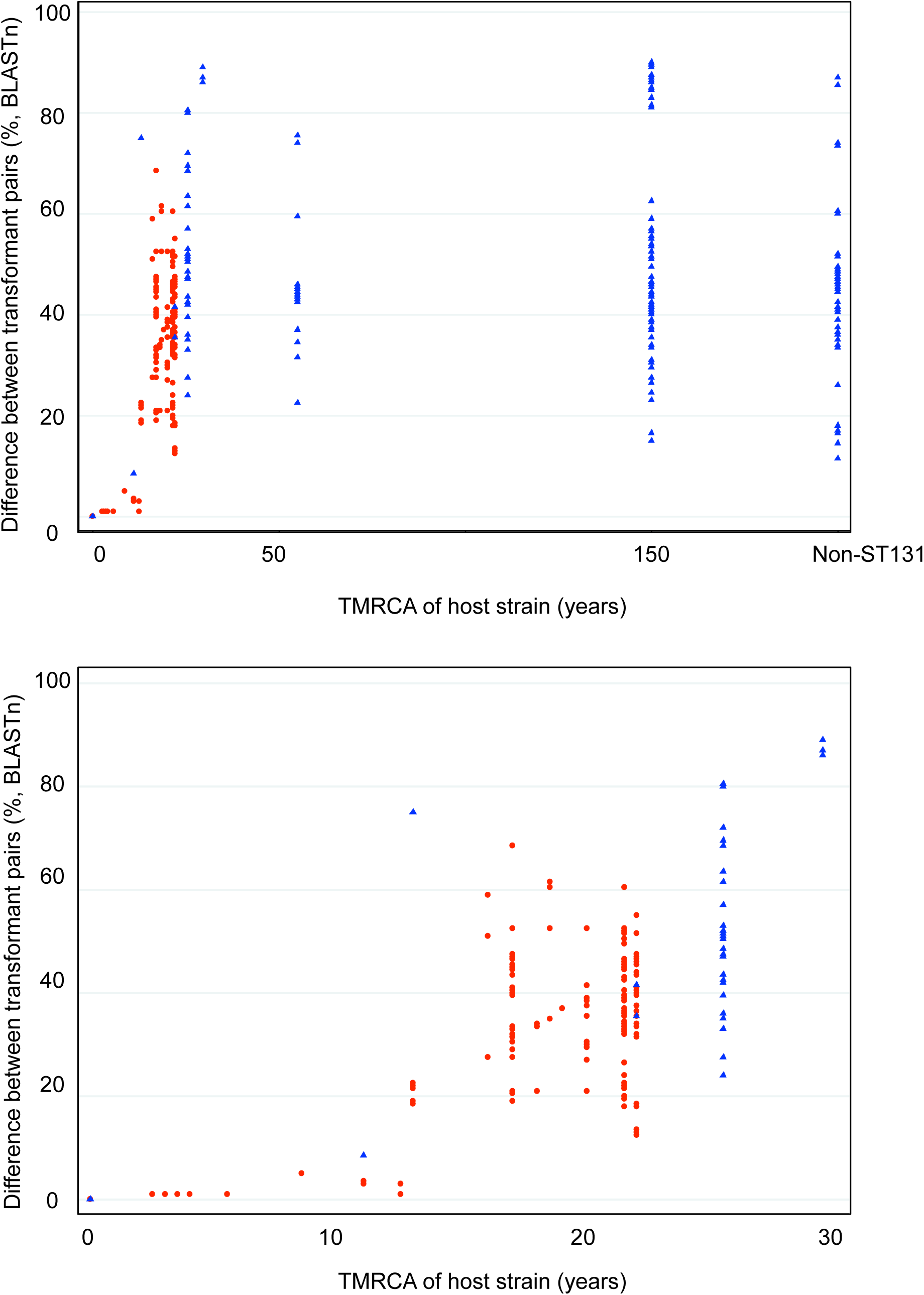
Mean pairwise percentage difference between all transformed plasmid sequence pairs plotted against time to most recent common ancestor (TMRCA) for the two strains hosting the respective transformant plasmids. Red circles indicate pairs where both strains are in C2; blue triangles indicate pairs where one or both strains are outside of C2. Lower panel represents the same data but limited to strains with a TMRCA of less than 30 years.

Most transformant plasmids were IncF, except in strains 11B00320_T and la_7619_T. BLASTn-based comparisons revealed that the clade A *bla*_CTX-M-15_ Incl transformant (11B00320_T; isolated in Mae Sot, Thai-Myanmar border) was circulating in a limited fashion (Fig. 5), but with substantial sequence homology to the other two clade A CTX-M-15-positive isolates (JJ2591, Minneapolis, USA and AZ779845, Spain). Although we did not have transformants or specific plasmid sequences for these, the *bla*_CTX-M-15_-containing contig assembled for JJ2591 was 88,693bp long and very similar to the 11B00320_T assembly, whereas the AZ779845 *bla*_CTX-M-15_-containing contig was 32,228bp long and likewise highly similar in structure (Fig. 5). These data suggest that an IncI-CTX-M-15 plasmid is responsible for sporadic, horizontal introductions of *bla*_CTX-M-15_ into ST131 with a wide geographic distribution.

**Figure 5.**
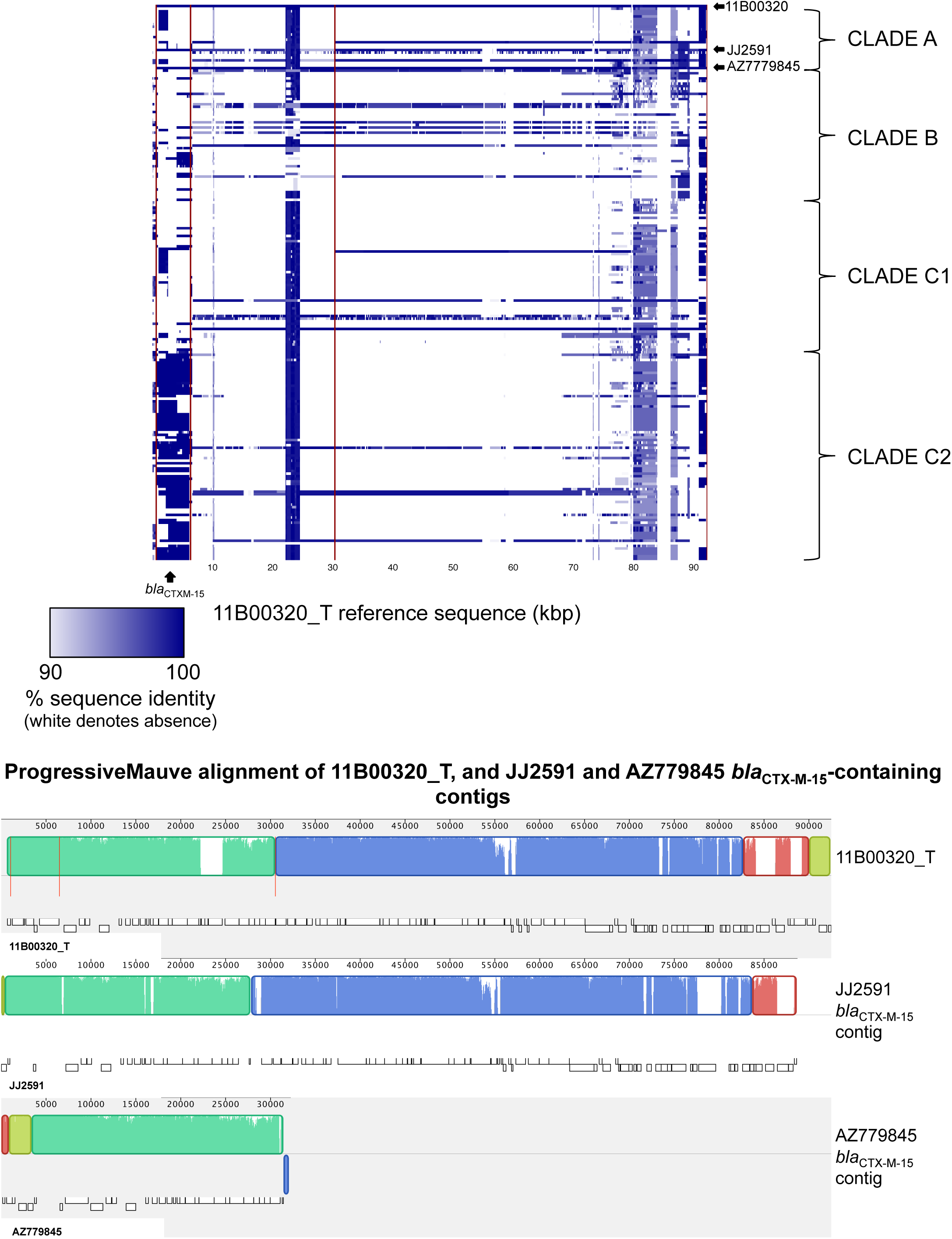
Top panel: BLASTn-based comparison across the ST131 dataset, using the *bla*_CTX-M-15_–containing 11B00320_T as a reference. Color represents degree of presence/absence of corresponding to the 11B00320_T sequence on an isolate-by-isolate basis per row. Rows/isolates are arranged as in the Fig. 1 phylogeny. Lower panel: ProgressiveMauve alignment of 11B00320_T, and the CTX-M-15 containing contigs for two other isolates in clade A. Alignments with substantial homology are represented as similarly colored blocks (“locally collinear blocks”); white regions within these blocks represent low homology. Vertical, red lines represent contig breaks.

Genetic comparisons amongst the *bla*_CTX-M-14/14-like_ transformant plasmids revealed that three shared strikingly similar genetic structures, two of which (uk_8A9B_T, Oxford, UK and cam_1071_T, Siem Reap, Cambodia) were identified in clade A, in host strains with a TMRCA within the last 15 years, and one in clade C1 (la_5108_T, Vientiane, Laos) (Fig. 6). BLASTn-based comparisons across all 215 ST131 sequences demonstrated that many isolates in clades A (predominantly sub-cluster [i]) and C1 contained highly similar genetic sequences, as did small numbers of isolates in clade C2. One of these, a *bla*_CTX-M-15_ transformant plasmid in clade C2, 11B01979_T (isolated in Mae Sot, Thai-Myanmar border), also showed significant homology to uk_8A9B_T, cam_1071_T and la_5108_T (Fig. 6); suggesting that both *bla*_CTX-M-14_ and *bla*_CTX-M-15_ variants can be accommodated on the same plasmid background.

**Figure 6.**
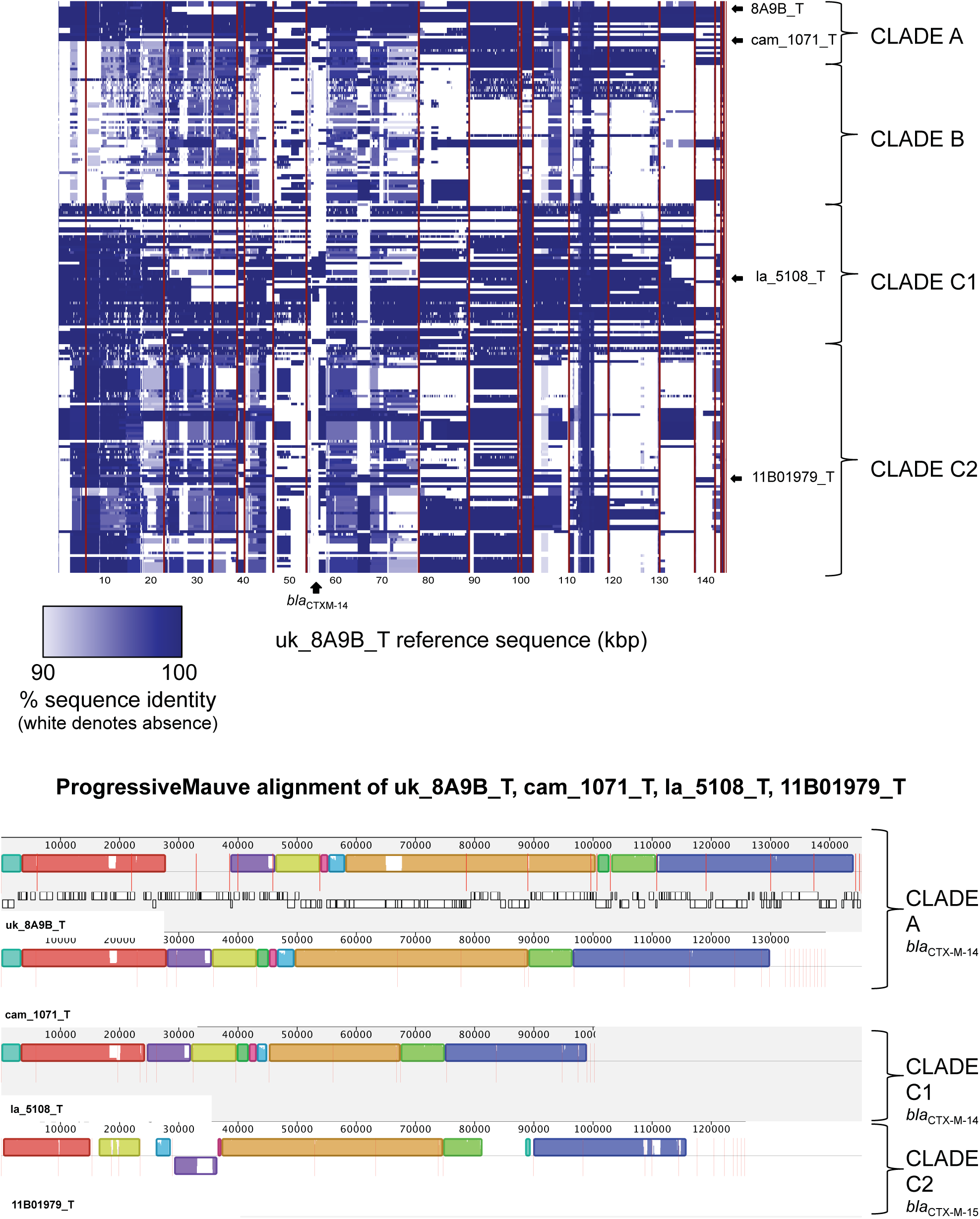
Top panel: BLASTn-based comparisons across the ST131 dataset, using the *bla*_CTX-M-14_–containing uk_8A9B_T as a reference. Color represents degree of presence/absence of corresponding to the uk_8A9B_T sequence on an isolate-by-isolate basis per row. Rows/isolates are arranged as in the Fig. 1 phylogeny. Lower panel: ProgressiveMauve alignment of uk_8A9B_T, caml071_T, la_5108_T and 11B01979_T, with the latter three ordered using uk_8A9B_T as a reference, and contig boundaries represented as vertical red lines. Alignments with substantial homology are represented as colored blocks (“locally collinear blocks”); white regions within these blocks represent low homology. Vertical, red lines represent contig breaks.

The isolates containing *bla*_CTX-M-55_ and *bla*_CTX-M-24_ (one SNV derivatives of *bla*_CTX-M-15_ and *bla*_CTX-M-14_, respectively) apparently resulted from discrete plasmid acquisition and/or *bla*_CTX-M_ transposition events within ST131 (Supplementary Figure S4). These were not therefore shown to represent *bla*_CTX-M_ evolution within established *bla*_CTX-M-15_ or *bla*_CTX-M-14_ plasmid backgrounds.

Nineteen of 20 *bla*_CTX-M-15_ transformant plasmids from clade C2 contained an FII_AY458016-like replicon, supporting the association of IncFII_AY458016 with *bla*_CTX-M-15_. When 17/20 IncFII_AY458016-containing transformant plasmids from clade C2 were compared with each other a significant degree of homology was evident (Fig. 7; excluding 8A16G_T, 11B01979_T, 19B19L_T – see methods). However, only eight coding sequences were shared with 100% nucleotide similarity, including: *bla*_CTX-M-15_, *bla*_oxA-1_, *aac(6’)-Ib-cr,* a glucose-l-phosphatase-like-enzyme, a CAAX amino terminal protease self-immunity protein, a hypothetical phage protein, and *apemI/pemK* plasmid addiction system. This lack of gene conservation suggests that significant genetic exchange and rearrangement occurs amongst these plasmids as they evolve within the sub-clade.

**Figure 7.**
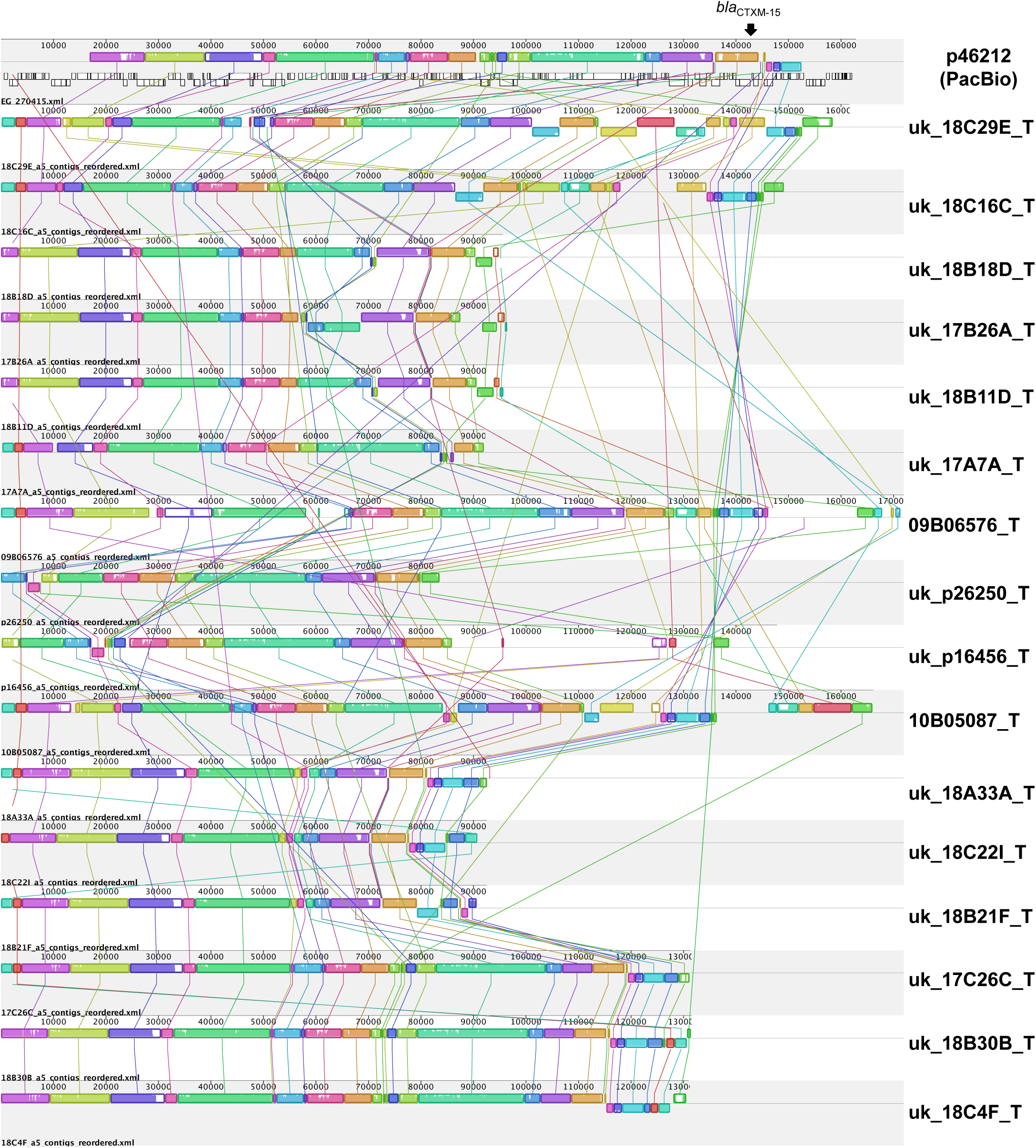
ProgressiveMauve alignment of assembled contigs, ordered with respect to pP46212, for 17 *bla*_CTX-M-15_ FII plasmids derived from sequenced transformants, all in clade C2/H30Rx. Plasmids are ordered with respect to the position of their host strains in the main phylogeny (except pP46212; Fig. 1). Alignments with substantial homology are represented as colored blocks (“locally collinear blocks”) and are linked with colored lines; white regions within these blocks represent low homology.

## DISCUSSION

Our WGS analysis of the largest and most diverse collection of ST131 isolates to date (n = 215) demonstrates conclusively that the global emergence of drug-resistant clades (*H*30, *H*30Rx) occurred approximately 25 years ago, most likely in a North American context, and consistent with strong selection pressure exerted by the widespread introduction and use of fluoroquinolones and extended-spectrum cephalosporins. Although members of each ST131 clade have dispersed globally, within specific geographic regions smaller clonal ST131 outbreaks occur at all genetic levels (gene, flanking context, plasmid, and host strain), indicating that both horizontal gene transfer and clonal expansion have contributed to the global dissemination of this sequence type. The estimated molecular evolutionary rate of ST131 (1.00 mutation per genome per year) is similar to previous estimates from ST131 [21] and the species overall [22], strongly suggesting that ST131’s epidemiological success is not due to a higher-than-average mutation rate.

Our study shows that the apparent persistence of particular *bla*_CTX-M_ variants within specific ST131 clades is due to diverse mechanisms. These include (i) acquisition of a *bla*_CTX-M_-containing plasmid by a specific host strain sub-cluster, followed by evolution and spread across geographic regions (e.g. clade A *bla*_CTX-M-14_ sub-cluster [i], Figs. 6 and Supplementary Figure S2); (ii) multiple discrete acquisition events involving *bla*_CTX-M_-containing plasmids (e.g. *bla*_CTX-M-55_, *bla*_CTX-M-24_; Supplementary Figure S4); (iii) horizontal transfer of common plasmid structures across clades (e.g. the IncI *bla*_CTX-M-15_ plasmid, Fig. 5); and (iv) chromosomal integration of *bla*_CTX-M_ and evolution by descent (e.g. *bla*_CTX-M-15_, Fig. 2; *bla*_CTX-M-14_). Despite this high degree of genetic plasticity, we also found clear structuring of *bla*_CTX-M_ variants and plasmid content, with the near-complete absence of *bla*_CTX-M_ in clade B, and associations of *bla*_CTX-M-14/14-like_ variants with clade A and clade C1/*H*30R, of *bla*_CTX-M-15_ with clade C2/*H*30Rx, and of specific combinations of IncF replicons with certain clades. This supports the hypothesis that some plasmid replicons are acquired and persist stably within clades. Although the evolutionary dynamics of plasmid-host combinations remain to be clearly elucidated, co-evolution of host and plasmid in the case of C2/*H*30Rx appear to have ameliorated costs to the host and facilitated persistence of the replicon[23, 24], with on-going conjugative exchange of genetic material. The relative contribution of changing environmental influences on this co-evolution is unclear; it may also be affected by a host-plasmid “arms race” in a micro-evolutionary version of the “Red Queen Hypothesis” (antagonistic co-evolution)[25, 26].

The almost ubiquitous presence of *bla*_CTX-M-15_ in clade C2/*H*30Rx is most striking, and is strongly associated with the acquisition of an IncFII_AY458016-like replicon. Previous smaller studies have found that *bla*_CTX-M-15_ is frequently part of a 2,971bp IS*Ecp1*-*bla*_CTX-M-15_-ORF477 transposition unit, with IS*Ecp1* located 48bp downstream of the IS*Ecp1* IRR-R, and that this is commonly nested within a Tn*2*-like element (IS*SWil*) [27], One hypothesis is that an IncFII_AY458016 ancestral plasmid was acquired by a fluoroquinolone-resistant C1 host strain approximately 25 years ago, and subsequently incorporated one of these *bla*_CTX-M-15_ transposition units. In response to the widespread clinical use of third-generation cephalosporins and fluoroquinolones the C2/*H*30Rx clade has expanded, and within it *bla*_CTX-M-15_ has been mobilized through further transposition events (e.g. to the chromosome) and rearrangement/recombination amongst IncFII-like plasmids, much of this associated with IS*26* [28] (Fig. 3). The persistence of the IncFII_AY458016-like replicon in C2 may be attributable, at least in part, to its association with aplasmid addiction system (*pemI/pemK*) [29], whereas its ongoing evolution is potentially linked to the concomitant presence of FIA/FIB replicons on *bla*_CTX-M-15_ plasmids (Supplementary Figure S3)[30], Alternative hypotheses – either of multiple, C2-restricted acquisitions of different *bla*_CTX-M-15_ -containing FII_AY458016-like plasmids, or of recurrent IS*Ecp1*-*bla*_CTX-M-15_**-**ORF477 unit acquisitions – seem less likely, given that (i) there are no geographic or major genotypic distinctions between clades C1 and C2 to explain why this would occur, (ii) there is a degree of homology in the flanking contexts around the gene throughout the clade, and (iii) flanking context/transformed plasmid structures also appear to be consistent within C2 sub-clusters.

Our novel comparison of transformed, sequenced plasmids demonstrates, however, that a substantial degree of similarity can exist amongst *bla*_CTX-M_ plasmids found in different clades and STs. This confirms that between-clade/ST transfer of these resistance plasmids occurs, and that care is needed when inferring plasmid evolution by descent (Fig. 4). Plasmid similarity across geography in the context of host strain phylogenetic clustering and homology in regions flanking *bla*_CTX-M_ (as demonstrated here) is much more likely to represent plasmid acquisition and evolution by descent rather than multiple acquisition events, but still needs to be interpreted with caution, as it may, for example, represent exposure to a common, global, plasmid reservoir.

The main study limitation is the inability with short-read sequencing and limited transformant sequencing to assess fully the flanking regions and plasmid structures across the entire dataset. In particular, the BLASTn-based heatmaps across the wider dataset represent not genetic contiguity of plasmid structures within isolates as such, but instead overall plasmid sequence presence/absence. Similarly, results from *de novo* assemblies of these short-read data also must be interpreted cautiously, as these assembly methods are known to increase the number of SNVs when compared with mapping-based approaches, and may result in misinterpretations of genetic structures, particularly repetitive regions [31], Further, again relating to the limitations of short-read data, the transformant plasmid sequences comprise multiple contigs, precluding certainty as to the plasmids’ exact structure. Wider use of long-read sequencing (e.g. PacBio) could help resolve this in future studies. Many of our *H*30Rx/C2 clade transformant plasmids were from a single UK center; however, the genetic flanking contexts identified here have also been found in plasmid sequences from other national and international locations [27, 32-34], suggesting that these are dispersed more widely and that our results are likely generalizable.

In summary, our analysis strongly suggests that the emergence of the C2/*H*30Rx clade within ST131 has been driven by the acquisition of a specific FII plasmid, which has subsequently undergone major genetic restructuring within its globally dispersing bacterial host. The initial acquisition event occurred approximately 25 years ago, possibly associated with the widespread introduction of third-generation cephalosporins and fluoroquinolones, which would have exerted significant selection pressure for persistence of chromosomal fluoroquinolone mutations and presence of *bla*_CTX-M_. Sporadic gain/loss events of other, non-FII *bla*_CTX-M-15_ plasmids have also occurred, but have not dominated. Similar processes may be driving the more recent emergence of sub-lineages of ST131 with *bla*_CTX-M-14_ and *bla*_CTX-M-27_, as described in Japan (18), although for *bla*_CTX-M-14_, these appear to have occurred on at least two occasions (clades A and C1/*H*30R; Fig. 1). This study highlights the global imperative to reduce antimicrobial selection pressures; the capacity of these resistance plasmids for genetic re-assortment; the important role of certain insertion sequences, such as IS*26*, in facilitating horizontal mobility of resistance determinants; and the possibility of targeting specific replicons in an attempt to limit the spread of important resistance gene mechanisms.

## METHODS

### Sample collection, sequencing and sequence read processing

Isolates were obtained from wider collections held in several centers: the Shoklo Malaria Research Unit, Mae Sot, Thailand; the Lao-Oxford-Mahosot Hospital Wellcome Trust Research Unit, Vientiane, People’s Democratic Republic of Laos; the Cambodia-Oxford Medical Research Unit, Angkor Hospital for Children, Siem Reap, Cambodia; the Microbiology Laboratory, Oxford University Hospitals NHS Trust, Oxford, UK. Strains were de-duplicated by individual prior to sequencing. In addition, seven isolates collected from clinical samples across Canada between 2006 and 2008, and one isolate recovered from poultry in 2006 were included. DNA was extracted as previously described [35]. Sequence data for the eight AstraZeneca strains had been generated from a series of isolates collected by International Health Management Associates, Inc, as part of a global resistance survey; that for the Price strains was as previously described [11]. Sequencing was performed using either the Illumina HiSeq or MiSeq (100 or 151bp paired-end reads [details for non-Price strains in Supplementary Table S1]). Correct sequence type was confirmed using BLASTn-based *in silico* MLST typing of *de novo* assembled WGS data [36].

Sequence data for all ST131 strains were mapped against the *E. coli* SE15 (ST131) reference (RefSeq: NC_013654) [17] and variants called using a validated in-house pipeline [37]. Alignments of core variable sites (base called in all sequences, excluding “N” or “-” calls) were reinserted into the reference to form an alignment of modified reference sequences.

*De novo* assemblies were generated using Velvet with the VelvetOptimizer wrapper (n=211), or A5-MiSeq [38], The latter was used in cases where the number of assembled bases was below the expected assembly size of 4-5.5Mb (n=4 [strains la_12107_3, can_70883, can_1731_01 and can_1070] in which median optimized assembly size with Velvet was 16,004 bases, and median number of contigs only six). Using A5-Miseq, assemblies for these four strains were generated with an appropriate median size of 5,143,908 bp and 269 contigs.

### Identification/characterization of *bla*_CTX-M_ and genetic context, *gyrA* mutations and *fimH* typing

BLASTn of *de novo* assemblies was used to identify: (i) *bla*_CTX-M_ presence and variant (in-house reference gene database) [35]; (ii) genetic context for *bla*_CTX-M_, by extracting and annotating contigs containing *bla*_CTX-M_ variants using PROKKA and ISFinder (manual annotation) [39, 40]; (iii) chromosomal *gyrA* mutations in the quinolone-resistance determining region known to be responsible for conferring most resistance to fluoroquinolones; (iv) *fimH* presence and variant [41]; and (v) Inc type using the downloaded PlasmidFinder [42] and pMLST databases (available at http://pubmlst.org/plasmid/) [43], Genetic contexts for *bla*_CTX-M_ were classified as chromosomal if annotations for regions flanking *bla*_CTX-M_ were found to be consistently chromosomal in other *E. coli* strains in GenBank, and plasmid if these were associated specifically with plasmids (e.g. *tra* genes); otherwise, they were classified as unknown. IncFII_AY458016-like sequences were extracted, aligned and visually inspected to confirm variant types using Geneious (Version 7.1.9; http://geneious.com) [44].

### ST131 chromosomal phylogenetic comparisons using ClonalFrame, BEAST and BASTA

ExPEC are recombinogenic, and contain recombination hotspots with higher than average recombination rates [45]. Recombination can obscure the true phylogenetic signal, and we therefore initially analyzed the alignment of sequences with ClonalFrame [46] to identify recombinant regions; any SNVs within these regions were excluded.

Using this modified alignment, mutation rate estimates across ST131 and atime-scaled phylogeny were calculated in BEAST [47], The model parameters were: (i) a generalized time-reversible nucleotide substitution model, (ii) four relative rates of mutation across sites, allowing for all sites to be subject to mutation (i.e. the proportion of invariant sites fixed at 0%), (iii) a strict molecular clock estimating a uniform evolutionary rate across all branches of the tree, and (iv) a constant population size. Triplicate runs with 30 million iterations were performed, with 10% discounted as burn-in. Run convergence and mixing was assessed by inspecting the run log files in Tracer v1.5 [48]; adequate convergence of run statistics and mixing for each run and effective sample sizes (ESS) for all parameters greater than 200 were required for an analysis to be considered adequate, in line with recommendations in the BEAST tutorials on the developers’ website (http://beast.bio.ed.ac.uk). We explored the application of several other models in BEAST incorporating the relaxed-clock and variable population growth (exponential, logistic and Bayesian skyride), but these either failed to converge, showed poor mixing, or had effective sample size estimates (ESS) of <200, and were therefore not considered robust.

We used the phylogeographic method BASTA [18] in the Bayesian phylogenetic package BEAST 2.2.1 [49] to infer patterns and rates of migration between geographical regions from the genome alignment, collection dates, and sampling locations. Initially, we grouped samples into three discrete locations: North America, South-East Asia, and Europe, and disregarded samples from South America and Australasia because of the small sample numbers. Due to the non-random sampling scheme, we only estimated a single effective population size, equal for all locations, and a symmetric migration rate matrix. The analysis was run for 10^8^ Monte Carlo Markov Chain (MCMC) steps. We subsequently re-ran the analysis including a fourth, unsampled deme, using the same model parameters, to determine whether this altered the outcome.

### Plasmid transformations, sequencing and analyses

Plasmid transformants were generated from 30 strains chosen on the basis of tree topology and association with CTX-M variants, aiming to transform at least one plasmid from each of the major CTX-M variant clusters. Two *bla*_CTX-M_ containing plasmids from non-ST131 *E. coli* (one ST617/*bla*_CTX-M-15_, one ST405/*bla*_CTX-M-55_) were also transformed and sequenced as an external comparison.

Plasmid DNA was extracted from sub-cultures of frozen stock grown overnight on blood agar, followed by selective culture of a single colony in Luria-Bertani (LB) broth with ceftriaxone at lμg/mL. DNA extraction was performed using the Qiagen plasmid mini-kit (Qiagen, Venlo, Netherlands), in accordance with the manufacturer’s instructions, with the addition of Glycoblue™ co-precipitate (Life Technologies, Carlsbad, USA) to the DNA eluates prior to isopropanol precipitation to enable better visualization of the DNA pellet. Plasmid DNA was re-dissolved in distilled water and then typically electroporated on the same day, or stored in the fridge prior to electroporation within 24 hours.

Commercially prepared DH10B *E. coli* (ElectroMAX™ DH10B™ Cells; Invitrogen/Life Technologies, Carlsbad, California, USA) were used as the recipient cell strain for plasmid electroporation, because of their high transformation efficiencies and the fact that the strain has been fully sequenced (NCBI RefSeq: NC_010473.1) [50], Briefly, 2μ1 of plasmid DNA (extracted as above) were mixed with 20μ1 of electrocompetent cells in a small Eppendorf tube on ice. The mix was then pipetted into a pre-chilled 0.2cm electroporation cuvette (Bio-Rad, Hercules, California, USA), placed in the MicroPulser™ electroporator, and an electric shock was applied (Ec2 settings; typically 2.5 kV applied for less than 5msec). The shocked cells were immediately suspended in 9mls of pre-warmed SOC medium (Super Optimal Broth with Catabolite repression) in a clean Eppendorf tube, and incubated at 37°C, whilst being shaken at 220rpm, for one hour. One hundred microliters of the transformant cell suspension was then plated onto pre-warmed LB agar plates (Becton Dickinson, Franklin Lakes, New Jersey, USA), infused with ceftriaxone (1μg/mL). pUC19 DNA was provided with the purchased cells and was used as a positive control for the success of transformation. Selective agars were incubated with known positive and negative control strains with each set of transformations.

Sequencing was performed on the IlluminaHiSeq or MiSeq generating 150 or 300-base paired-end reads (Supplementary Table S2). Sequencing reads from the isolate from which the transformed plasmid had been obtained were mapped back to the transformed plasmid assembly in order to ascertain the reliability of the assembly in each case. Reads were assembled using A5/A5-MiSeq [38], and assembled contigs annotated with PROKKA [39]. The median plasmid assembly size was 122,786 (range: 72,449-171,919), with a median of 22 contigs in each assembly (range: 1-33). Using longer reads (300bp; MiSeq platform) resulted in a significantly smaller number of contigs per assembly (median 17 versus 25; ranksum p=0.003). Mapping was used to assess the reliability of our plasmid constructs and reflected the content present in each transformed and assembled resistance plasmid, with the exception of 8A16G_T.

A single strain (P46212) from the dataset was also sequenced using long-read technology (PacBio); the CTX-M-15 plasmid (pP46212) from this strain was assembled into a single, circularized contig as described [51].

Plasmid content across the dataset was investigated in a number of ways. Firstly, the transformant plasmid sequences generated were used as references, against which BLASTn-based comparisons for degree of presence/absence were made for the whole dataset. We used default BLASTn settings, with the respective reference divided into 100bp bins. Secondly, pairwise comparisons between each set of transformant assemblies were undertaken by identifying the extent of shared homology of sequences across a number of subsets representing different groupings on the main host tree, again using BLASTn. For each pair, two percentage-similarity statistics were generated, taking each member of the pair as a reference in turn, to account for differences in length, and using default BLASTn settings. The mean percentage pairwise divergence was then plotted against the time to most recent common ancestor (TMRCA) of the two host strains (derived from the time-scaled tree). Thirdly, for visualization, plasmid sequences were compared using ProgressiveMauve [52], with assembled contigs reordered with respect to the pP46212 PacBio-generated CTX-M-15 plasmid reference, using the “Move contigs” tool. For this three transformants were excluded: 8A16G_T because of issues surrounding the assembly, 11B01979_T because it was virtually identical to *bla*_CTX-M-14_ plasmid transformants in clade A, and 19B19L_T because it lacked an FII replicon. Finally, annotated, nucleotide sequences across transformant groups of interest were clustered using CD-Hit [53] [-c 1.0 -n 5 -d 0 -g 1],to identify whether any coding sequences were shared, and whether there might be any biological significance associated with these on the basis of their annotations.

Sequencing data for the new isolates sequenced for this study have been deposited in the NCBI short read archive (BioProject number: PRJNA297860, 108 ST131 sequences and 30 *bla*_CTX-M_ plasmid transformants).

## ACKNOWLEDGEMENTS

We are grateful to the patients and staff at the healthcare, microbiology laboratory and research units contributing isolates to this study, including Prof Nicholas Day of the Mahidol-Oxford Tropical Medicine Research Unit, Bangkok, Thailand; Prof Paul Newton and Dr David Dance of the Lao-Oxford-Mahosot Hospital-Wellcome Trust Research Unit, Vientiane, Laos; the Ped Study Team and Microbiology Laboratory at Patan Hospital, Kathmandu, Nepal; and Prof Francois Nosten of the Shoklo Malaria Research Unit, Mae Sot, Thailand. We thank Prof Peter Donnelly and the staff at the Sequencing Center, Wellcome Trust Center for Human Genetics, Oxford, UK, for their sequencing work, and Dr Laura Matseje of the Public Health Agency of Canada for sharing her laboratory protocol for plasmid transformation. We are grateful to Prof Johann Pitout, Prof Nicholas Day, Dr Amy Mathers and Dr Chris Parry for their critical review of the draft manuscript.

## SUPPLEMENTARY FILE LEGENDS

Supplementary Table S1. Details of new ST131 strains included in the analysis.

Supplementary Figure S1. Genetic contexts of clade A-associated *bla*_CTX-M-14/14-like_ variants. Many contexts are limited by the extent of the assembled region around the *bla*_CTX-M-14/14-like_ gene (marked with “X”). For all aligned, similarly colored regions, sequence homology is preserved; curly brackets cluster those isolates with identical flanking sequences. Flanking contexts not shown for isolates with known chromosomal integration, or for *bla*_CTX-M_ negative/non-*bla*_CTX-M-14/14-like_ isolates in the sub-clusters. Coloring of isolate names reflects geographic locations (blue = North America, red = Europe, green = South-East Asia, Yellow = Australasia).

Supplementary Figure S2. Genetic contexts of clade C1-associated *bla*_CTX-M-14/14-like_ variants. Many contexts are limited by the extent of the assembled region around the *bla*_CTX-M-15_ gene (marked with “X”). For all aligned, similarly colored regions, sequence homology is preserved; curly brackets cluster those isolates with identical flanking sequences. Flanking contexts not shown for isolates with known chromosomal integration, or for *bla*_CTX-M_ negative/non-*bla*_CTX-M-14/14-like_ isolates in the sub-cluster, (blue = North America, red = Europe, green = South-East Asia, Yellow = Australasia).

Supplementary Figure S3. Inc types identified in whole isolate sequencing data, plotted with respect to ST131 host strain phylogeny. Blast match (%) denotes a composite score of percentage matched length and percentage homology to reference Inc sequence, with highest percentage score/contig hit represented. Matches < 80% were excluded. Reference Inc sequences were downloaded from the PlasmidFinder database; those that were present (Blast match ≥ 80%) in at least one isolate are represented on the x-axis.

Supplementary Table S2. Details of transformed *bla*_CTX-M_ plasmid sequences.

Supplementary Figure S4. BLASTn-based comparisons across the ST131 dataset, using la_12107–3 and la_5220–3_T as references. Color represents degree of presence/absence of corresponding reference sequence on an isolate-by-isolate basis per row. Rows/isolates arranged as in the Fig. 1 phylogeny.

## REFERENCES

1. Rogers BA, Sidjabat HE, Paterson DL: *Escherichia coli* O25b-ST131: a pandemic, multiresistant, community-associated strain. J Antimicrob Chemother 2011, 66:1–14.

2. Woodford N, Turton JF, Livermore DM: Multiresistant Gram-negative bacteria: the role of high-risk clones in the dissemination of antibiotic resistance. FEMS Microbiol Rev 2011, 35:736–755.

3. Coque TM, Novais A, Carattoli A, Poirel L, Pitout J, Peixe L, Baquero F, Canton R, Nordmann P: Dissemination of clonally related *Escherichia coli* strains expressing extended-spectrum beta-lactamase CTX-M-15. Emerg Infect Dis 2008, 14:195–200.

4. Nicolas-Chanoine MH, Blanco J, Leflon-Guibout V, Demarty R, Alonso MP, Canica MM, Park YJ, Lavigne JP, Pitout J, Johnson JR: Intercontinental emergence of *Escherichia coli* clone O25:H4-ST131 producing CTX-M-15. J Antimicrob Chemother 2008, 61:273–281.

5. Mathers AJ, Peirano G, Pitout JD: *Escherichia coli ST131*: The quintessential example of an international multiresistant high-risk clone. Adv Appl Microbiol 2015, 90:109–154.

6. Nicolas-Chanoine MH, Bertrand X, Madec JY: *Escherichia coli* ST131, an intriguing clonal group. Clin Microbiol Rev 2014, 27:543–574.

7. Brisse S, Diancourt L, Laouenan C, Vigan M, Caro V, Arlet G, Drieux L, Leflon-Guibout V, Mentre F, Jarlier V, et al: Phylogenetic distribution of CTX-M- and non-extended-spectrum-beta-lactamase-producing *Escherichia coli* isolates: group B2 isolates, except clone ST131, rarely produce CTX-M enzymes. J Clin Microbiol 2012, 50:2974–2981.

8. Canton R, Gonzalez-Alba JM, Galan JC: CTX-M Enzymes: Origin and Diffusion. Front Microbiol 2012, 3:110.

9. Naseer U, Sundsfjord A: The CTX-M conundrum: dissemination of plasmids and *Escherichia coli* clones. Microb Drug Resist 2011, 17:83–97.

10. Novais A, Pires J, Ferreira H, Costa L, Montenegro C, Vuotto C, Donelli G, Coque TM, Peixe L: Characterization of globally spread *Escherichia coli* ST131 isolates (1991 to 2010). Antimicrob Agents Chemother 2012, 56:3973–683 3976.

11. Price LB, Johnson JR, Aziz M, Clabots C, Johnston B, Tchesnokova V, Nordstrom L, Billig M, Chattopadhyay S, Stegger M, et al: The epidemic of extended-spectrum-beta-lactamase-producing *Escherichia coli* ST131 is driven by a single highly pathogenic subclone, H30-Rx. MBio 2013, 4:e00377–00313.

12. Kim J, Bae IK, Jeong SH, Chang CL, Lee CH, Lee K: Characterization of IncF plasmids carrying the *bla*_CTX-M-14_ gene in clinical isolates of *Escherichia coli* from Korea. J Antimicrob Chemother 2011, 66:1263–1268.

13. Rodriguez I, Thomas K, Van Essen A, Schink AK, Day M, Chattaway M, Wu G, Mevius D, Helmuth R, Guerra B, consortium S-E: Chromosomal location of *bla*_CTX-M_ genes in clinical isolates of *Escherichia coli* from Germany, The Netherlands and the UK. Int J Antimicrob Agents 2014, 43:553–557.

14. Petty NK, Ben Zakour NL, Stanton-Cook M, Skippington E, Totsika M, Forde BM, Phan MD, Gomes Moriel D, Peters KM, Davies M, etal: Global dissemination of a multidrug resistant *Escherichia coli* clone. Proc Natl Acad Sci U S A 2014, 111:5694–5699.

15. Woerther PL, Burdet C, Chachaty E, Andremont A: Trends in human fecal carriage of extended-spectrum beta-lactamases in the community: toward the globalization of CTX-M. Clin Microbiol Rev 2013, 26:744–758.

16. Jean SS, Hsueh PR: High burden of antimicrobial resistance in Asia. Int J Antimicrob Agents 2011, 37:291–295.

17. Toh H, Oshima K, Toyoda A, Ogura Y, Ooka T, Sasamoto H, Park SH, Iyoda S, Kurokawa K, Morita H, et al: Complete genome sequence of the wild-type commensal *Escherichia coli* strain SE15, belonging to phylogenetic group B2. J Bacteriol 2010, 192:1165–1166.

18. De Maio N, Wu CH, O’Reilly KM, Wilson D: New Routes to Phylogeography: A Bayesian Structured Coalescent Approximation. PLoS Genet 2015, 11:e1005421.

19. Ruiz J: Mechanisms of resistance to quinolones: target alterations, decreased accumulation and DNA gyrase protection. J Antimicrob Chemother 2003, 51:1109–1117.

20. Poirel L, Lartigue MF, Decousser JW, Nordmann P: IS*Ecp1*B-mediated transposition of *bla*_CTX-M_ in *Escherichia coli*. Antimicrob Agents Chemother 2005, 49:447–450.

21. Reeves PR, Liu B, Zhou Z, Li D, Guo D, Ren Y, Clabots C, Lan R, Johnson JR, Wang L: Rates of mutation and host transmission for an *Escherichia coli* clone over 3 years. PLoS One 2011, 6:e26907.

22. Ochman H: Neutral mutations and neutral substitutions in bacterial genomes. Mol Biol Evol 2003, 20:2091–2096.

23. Bahl MI, Hansen LH, Sorensen SJ: Persistence mechanisms of conjugative plasmids. Methods Mol Biol 2009, 532:73–102.

24. Harrison E, Guymer D, Spiers AJ, Paterson S, Brockhurst MA: Parallel Compensatory Evolution Stabilizes Plasmids across the Parasitism-Mutualism Continuum. Curr Biol 2015, 25:2034–2039.

25. Harrison E, Brockhurst MA: Plasmid-mediated horizontal gene transfer is a coevolutionary process. Trends Microbiol 2012, 20:262–267.

26. Brockhurst MA, Chapman T, King KC, Mank JE, Paterson S, Hurst GD: Running with the Red Queen: the role of biotic conflicts in evolution. Proc Biol Sci 2014, 281.

27. Partridge SR, Zong Z, Iredell JR: Recombination in IS*26* and Tn*2* in the evolution of multiresistance regions carrying *bla*_CTX-M-15_ on conjugative IncF plasmids from *Escherichia coli*. Antimicrob Agents Chemother 2011, 55:4971–4978.

28. He S, Hickman AB, Varani AM, Siguier P, Chandler M, Dekker JP, DydaF: Insertion Sequence IS*26* Reorganizes Plasmids in Clinically Isolated Multidrug-Resistant Bacteria by Replicative Transposition. MBio 2015, 6.

29. Carattoli A: Plasmids and the spread of resistance. Int J Med Microbiol 2013, 303:298–304.

30. Sykora P: Macroevolution of plasmids: a model for plasmid speciation. J Theor Biol 1992, 159:53–65.

31. Stoesser N: Applications of whole genome sequencing to understanding the mechanisms, evolution and transmission of antibiotic resistance in Escherichia coli and Klebsiella pneumoniae. Oxford, 2014.

32. Boyd DA, Tyler S, Christianson S, McGeer A, Muller MP, Willey BM, Bryce E, Gardam M, Nordmann P, Mulvey MR: Complete nucleotide sequence of a 92-749 kilobase plasmid harboring the CTX-M-15 extended-spectrum beta-lactamase involved in an outbreak in long-term-care facilities in Toronto, Canada. Antimicrob Agents Chemother 2004, 48:3758–3764.

33. Matsumura Y, Johnson JR, Yamamoto M, Nagao M, Tanaka M, Takakura S, Ichiyama S, Kyoto-Shiga Clinical Microbiology Study G, Kyoto-Shiga Clinical Microbiology Study G: CTX-M-27- and CTX-M-14-producing, ciprofloxacin-resistant *Escherichia coli* of the H30 subclonal group within ST131 drive a Japanese regional ESBL epidemic. J Antimicrob Chemother 2015, 70:1639–757 1649.

34. Woodford N, Carattoli A, Karisik E, Underwood A, Ellington MJ, Livermore DM: Complete nucleotide sequences of plasmids pEK204, pEK499, and pEK516, encoding CTX-M enzymes in three major *Escherichia coli* lineages from the United Kingdom, all belonging to the international O25:H4-ST131 clone. Antimicrob Agents Chemother 2009, 53:4472–4482.

35. Stoesser N, Batty EM, Eyre DW, Morgan M, Wyllie DH, Del Ojo Elias C, Johnson JR, Walker AS, Peto TE, Crook DW: Predicting antimicrobial susceptibilities for *Escherichia coli* and *Klebsiella pneumoniae* isolates using whole genomic sequence data. J Antimicrob Chemother 2013, 68:2234–2244.

36. Wirth T, Falush D, Lan R, Colles F, Mensa P, Wieler LH, Karch H, Reeves PR, Maiden MC, Ochman H, Achtman M: Sex and virulence in *Escherichia coli:* an evolutionary perspective. Mol Microbiol 2006, 60:1136–1151.

37. Eyre DW, Cule ML, Wilson DJ, Griffiths D, Vaughan A, O’Connor L, Ip CL, Golubchik T, Batty EM, Finney JM, et al: Diverse sources of *C. difficile* infection identified on whole-genome sequencing. N Engl J Med 2013, 369:1195–1205.

38. Coil D, Jospin G, Darling AE: A5-miseq: an updated pipeline to assemble microbial genomes from Illumina MiSeq data. Bioinformatics 2015, 31:587–589.

39. Seemann T: Prokka: rapid prokaryotic genome annotation. Bioinformatics 2014, 30:2068–2069.

40. Siguier P, Perochon J, Lestrade L, Mahillon J, Chandler M: ISfinder: the reference centre for bacterial insertion sequences. Nucleic Acids Res 2006, 34:D32–36.

41. Weissman SJ, Johnson JR, Tchesnokova V, Billig M, Dykhuizen D, Riddell K, Rogers P, Qin X, Butler-Wu S, Cookson BT, et al: High-resolution two-locus clonal typing of extraintestinal pathogenic *Escherichia coli*. Appl Environ Microbiol 2012, 78:1353–1360.

42. Carattoli A, Zankari E, Garcia-Femandez A, Voldby Larsen M, Lund O, Villa L, Moller Aarestrup F, Hasman H: In silico detection and typing of plasmids using PlasmidFinder and plasmid multilocus sequence typing. Antimicrob Agents Chemother 2014, 58:3895–3903.

43. Jolley KA, Maiden MC: BIGSdb: Scalable analysis ofbacterial genome variation at the population level. BMC Bioinformatics 2010, 11:595.

44. Kearse MM. R.; Wilson, A.; Stones-Havas, S.; Cheung, M.; Sturrock, S.; Buxton, A.; Markowitz, S.; Duran, C.; Thierer, T.; Ashton, B.; Metjies, P.; Drummond A.: Geneious Basic: an integrated and extendable desktop software platform for the organization and analysis of sequence data. Bioinformatics 2012, 28:1647–796 1649.

45. Didelot X, Meric G, Falush D, Darling AE: Impact of homologous and non-homologous recombination in the genomic evolution of *Escherichia coli*. BMC Genomics 2012, 13:256.

46. Didelot X, Falush D: Inference of bacterial microevolution using multilocus sequence data. Genetics 2007, 175:1251–1266.

47. Drummond AJ, Suchard MA, Xie D, Rambaut A: Bayesian phylogenetics with BEAUti and theBEAST 1.7. Mol Biol Evol 2012, 29:1969–1973.

48. Tracer vl.6 [http://beast.bio.ed.ac.uk/Tracer]

49. Bouckaert R, Heled J, Kuhnert D, Vaughan T, Wu CH, Xie D, Suchard MA, Rambaut A, Drummond AJ: BEAST 2: a software platform for Bayesian evolutionary analysis. PLoS ComputBiol 2014, 10:e1003537.

50. Durfee T, Nelson R, Baldwin S, Plunkett G, 3rd, Burland V, Mau B, Petrosino JF, Qin X, Muzny DM, Ayele M, et al: The complete genome sequence of *Escherichia coli* DH10B: insights into the biology of a laboratory workhorse. J Bacteriol 2008, 190:2597–2606.

51. Sheppard AS. N.; Wilson, D.J.; Sebra, R.; Kasarskis, A.; Anson, L.W.; Giess, A.; Pankhurst, L.J.; Vaughan, A.; Grim, C.J.; Cox, H.L.; Yeh, A.J.; Modernising Medical Microbiology Informatics Group; Sifri, C.D.; Walker, A.S.; Peto, T.E.A.; Crook, D.W.; Mathers, A.J.: Rapid spread of a carbapenem resistance gene driven by multiple levels of genetic mobility. In submission to Nature Microbiology 2015.

52. Darling AE, Mau B, PemaNT: progressiveMauve: multiple genome alignment with gene gain, loss and rearrangement. PLoS One 2010, 5:e11147.

53. Li W, Godzik A: Cd-hit: a fast program for clustering and comparing large sets of protein or nucleotide sequences. Bioinformatics 2006, 22:1658–1659.

